# White matter fascicles and cortical microstructure predict reading-related responses in human ventral temporal cortex

**DOI:** 10.1101/2020.04.13.040063

**Authors:** Mareike Grotheer, Jason Yeatman, Kalanit Grill-Spector

**Affiliations:** Psychology Department, Stanford University, Stanford, CA 94305, USA; Graduate School of Education, Stanford University, Stanford, CA 94305, USA; Division of Developmental-Behavioral Pediatrics, Stanford University School of Medicine, Stanford, CA 94305, USA

**Keywords:** visual word form area, white matter, gray matter, reading, vision

## Abstract

Reading-related responses in the lateral ventral temporal cortex (VTC) show a consistent spatial layout across individuals, which is puzzling, since reading skills are acquired during childhood. Here, we tested the hypothesis that white matter fascicles and gray matter microstructure predict the location of reading-related responses in lateral VTC. We obtained functional (fMRI), diffusion (dMRI), and quantitative (qMRI) magnetic resonance imaging data in 30 adults. fMRI was used to map reading-related responses by contrasting responses in a reading task with those in adding and color tasks; dMRI was used to identify the brain’s fascicles and to map their endpoints density in lateral VTC; qMRI was used to measure proton relaxation time (T_1_), which depends on cortical tissue microstructure. We fit linear models that predict reading-related responses in lateral VTC from endpoint density and T_1_ and used leave-one-subject-out cross-validation to assess prediction accuracy. Using a subset of our participants (N=10, feature selection set), we find that i) endpoint density of the arcuate fasciculus (AF), inferior longitudinal fasciculus (ILF), and vertical occipital fasciculus (VOF) are significant predictors of reading-related responses, and ii) cortical T_1_ of lateral VTC further improves the predictions of the fascicle model. Next, in the remaining 20 participants (validation set), we showed that a linear model that includes T_1_, AF, ILF and VOF significantly predicts i) the map of reading-related responses across lateral VTC and ii) the location of the visual word form area, a region critical for reading. Overall, our data-driven approach reveals that the AF, ILF, VOF and cortical microstructure have a consistent spatial relationship with an individual’s reading-related responses in lateral VTC.

**Highlights:** The ILF, AF, and VOF predict the spatial layout of reading-related responses in VTC

Gray matter microstructure improves the prediction of reading-related responses

Fascicles and gray matter structure together predict the location of the VWFA

## 1. Introduction

The interplay between brain structure and function in concerting human cognition has spiked considerable interest in recent years. One of the most fascinating observations in this context is the consistent spatial arrangement of category-selective regions in ventral temporal cortex (VTC) across individuals (Glezer and Riesenhuber, 2013; Grill-Spector et al., 2017; Grill-Spector and Weiner, 2014; Kanwisher, 2010; Weiner et al., 2018; Weiner and Grill-Spector, 2010). VTC contains several visual category-selective regions that respond more strongly to their preferred category than others, including regions that are selective to faces (Kanwisher et al., 1997), scenes (Epstein and Kanwisher, 1998), bodies (Peelen and Downing, 2005), words (Cohen et al., 2000) and possibly numbers (Grotheer et al., 2016b, 2016a; Shum et al., 2013, but see Grotheer et al., 2018). These category-selective regions play a critical role in visual perception; for instance, a lesion of a category-selective region produces a specific inability to recognize items of that category (Gaillard et al., 2006; Konen et al., 2011; Rossion et al., 2003; Schiltz et al., 2006); similarly, electrical stimulation of a category-selective region specifically disrupts the perception of the respective category (Jonas et al., 2012; Julian et al., 2016; Parvizi et al., 2012; Rangarajan et al., 2014). Strikingly, recent research has shown that anatomical landmarks can predict the location of these category-selective regions in VTC. For example, the mid-fusiform sulcus predicts the location of face-selective regions in the fusiform gyrus (Weiner et al., 2014) and the intersection between the anterior lingual sulcus and the collateral sulcus predict the location of the place-selective region (Weiner et al., 2018). However, presently, there is an active debate regarding what factors drive this strikingly consistent spatial organization of VTC.

Several hypotheses have been put forward to explain the consistent localization of category-selective regions across individuals, including: i) genetics, which may innately determine the location of these regions (Abbasi et al., 2020; McKone et al., 2012; Polk et al., 2007), ii) foveal and peripheral eccentricity biases coupled with consistent viewing demands of certain stimuli (e.g., foveation on words during reading) (Behrmann and Plaut, 2015; Hasson et al., 2002; Malach et al., 2002), (iii) the underlying cortical microarchitecture, which is supported by the observation that different category-selective regions are located within different cytoarchitectonic regions of VTC (Gomez et al., 2017; Weiner et al., 2017) and iv) white matter connections, as category-selective regions are part of more extended brain networks and hence need to communicate with other regions across the brain (Haxby et al., 2000; Osher et al., 2016; Papagno et al., 2011; Saygin et al., 2016, 2012). Note that these hypotheses are not mutually exclusive as all of these factors together may constrain the location of functional regions (e.g., see Behrmann and Plaut, 2015a for a unifying framework).

Here, we primarily focus on the last hypothesis and evaluate if and how the location of major white matter fascicles relates to functional responses in VTC while participants engage in a reading task. We focus on the white matter connections of reading for two main reasons. First, given that reading is a learned skill, it is particularly puzzling what drives the consistent spatial layout of reading-related responses across individuals. Indeed, recent evidence shows that learning to read changes functional response in the lateral portion of VTC (lateral VTC) (Ben-Shachar et al., 2011; Cantlon et al., 2011), leading not only to changes in distributed responses (Nordt et al., 2019), but also to the emergence of a region selective for words (Dehaene-Lambertz et al., 2018; Dehaene et al., 2010). This region, which is often referred to as the visual word form area (VWFA, Cohen et al., 2000; Dehaene and Cohen, 2011), is located in the occipito-temporal sulcus (OTS) and critically involved in reading (Gaillard et al., 2006; Hirshorn et al., 2016). Second, recent research has provided evidence for the idea that white matter connections predict the location of category-selective regions involved in processing faces, words, places, and tools (Bi et al., 2015; Osher et al., 2016; Saygin et al., 2016, 2012), and, in particular, play a crucial role in constraining the spatial layout of reading-related responses in lateral VTC (Saygin et al., 2016). Specifically, Saygin and colleagues reported that pairwise white matter connections between cortical regions, the so called “white matter fingerprint”, at age 5 predicts functional responses in VTC to words at age 8. However, an open question remains as to which white matter fascicles actually constitute this “white matter fingerprint” and predict the spatial layout of reading-related responses in lateral VTC.

To address this gap in knowledge, we examined which white matter fascicles of the human brain predict the map of reading-related responses across lateral VTC as well as the location of the VWFA. White matter fascicles are the large-scale bundles that connect distant cortical regions. Several studies have identified the fascicles that connect to the VWFA (Bouhali et al., 2014; Grotheer et al., 2019; Lerma-Usabiaga et al., 2018; Yeatman et al., 2013) and those that contribute to the reading network as a whole (Grotheer et al., 2019). These fascicles include i) the arcuate fasciculus (AF), which connects the temporal and the frontal lobes (Catani et al., 2002), ii) the posterior arcuate fascicle (pAF), which connects the temporal and the parietal lobes (Weiner et al., 2016), iii) the inferior longitudinal fasciculus (ILF) which connects the occipital lobe with the anterior tip of the temporal lobe (Catani et al., 2002), iv) the inferior fronto-occipital fasciculus (IFOF), which connects the occipital and the frontal lobes (Catani et al., 2002) and v) the VOF, which connects the occipital and the parietal lobes (Takemura et al., 2016; Weiner et al., 2016; Yeatman et al., 2014b). However, no previous studies tested the hypothesis that these fascicles covary with the spatial layout of reading-related responses in lateral VTC. In other words, it is unknown if the endpoints of these fascicles in lateral VTC co-localize with the spatial map of reading-related response as well as the location of the VWFA. Critically, recent methodological innovations in white matter tractography, including constrained spherical deconvolution (CSD, Tournier et al., 2019, 2012) and anatomically-constrained tractography (ACT, Smith et al., 2012), now enable researchers to resolve white matter tracts close to the white matter / gray matter boundary, thereby allowing us to directly investigate if and how fascicles contribute to the consistent spatial layout of reading-related responses in lateral VTC.

While we will primarily focus on the role of white matter fascicles in driving the spatial layout of reading-related responses in lateral VTC, we also examined an additional hypothesis that has not been tested previously. That is, we asked whether differences in the gray matter microstructure across the cortical surface may also contribute to the consistent localization of category-selective regions in VTC. Recently developed quantitative MRI (qMRI) methods (Edwards et al., 2018; Lutti et al., 2014; Mezer et al., 2013) now enable assessing characteristics of the tissue microstructure of the gray matter *in vivo* (Gomez et al., 2017; Lutti et al., 2014; Natu et al., 2019; Weiskopf et al., 2013). QMRI measures proton relaxation time (T_1_), which differs between distinct category-selective regions (Gomez et al., 2017), changes during development (Gomez et al., 2017; Natu et al., 2019), and correlates with people’s performance in certain tasks, such as face processing (Gomez et al., 2017). As T_1_ depends on the local cortical microstructure (such as myelination, cell density, cell distribution, and iron concentration), which also covaries with the location of functional regions (Weiner et al., 2017), it is an open question whether T_1_ may be an additional factor predicting the spatial layout of reading-related responses in lateral VTC.

To address the hypothesis that white matter fascicles and gray matter microstructure contribute to the consistent functional organization of VTC, we used a mutimodal approach in which we obtained functional (fMRI), diffusion (dMRI) and quantitative MRI (qMRI) data in 30 adult participants. Using fMRI, we identified reading-related responses in lateral VTC by contrasting activations during a reading task with those elicited by adding and color tasks. These three tasks were performed on idenical visual stimuli (letter/number morphs) as in prior studies (**Fig. 1a**, Grotheer et al., 2019, 2018), allowing us to disentangle the impact of the performed task from that of the visual stimulus the task is being performed on. Using this contrast, we (1) mapped activations during reading across lateral VTC, and (2) identified a region in the mid occipito temporal sulcus (mOTS) that shows a significant preference for the reading task over the other tasks (T≥3, voxel level) (**Fig. 1b**). This functional region of interest (fROI) likely corresponds to the VWFA-2 (Grotheer et al., 2018; Lerma-Usabiaga et al., 2018; White et al., 2019). Using dMRI data, we generated a white matter connectome in each participant, automatically identified the white matter fascicles that connect to lateral VTC (**Fig. 1c**), and mapped the endpoint density of these fascicles to the cortical surface. Using qMRI, we meassured proton relaxation time (T_1_) in each particpant and mapped T_1_ across the cortical surface (**Fig. 1d**). To determine which are the informative features of the white matter anatomy that predict the spatial layout of reading-related responses in lateral VTC, we used data from 10 randomly selected participants (the feature selection set) and derived linear models relating fascicle endpoint densities to reading-related reponses in lateral VTC (**Fig. 1e**). Next, to test if gray matter properties improve the prediction of reading-related responses in lateral VTC compared to using only fascicle endpoints, we tested if adding the T_1_ value of each vertex improves the model. Finally, using independent data from 20 new subjects (the validation set), we tested how well a model that combines the most informative features of the white and gray matter anatomy predicts the map of reading-related responses across lateral VTC as well the location of the VWFA-2.

**Fig. 1.**
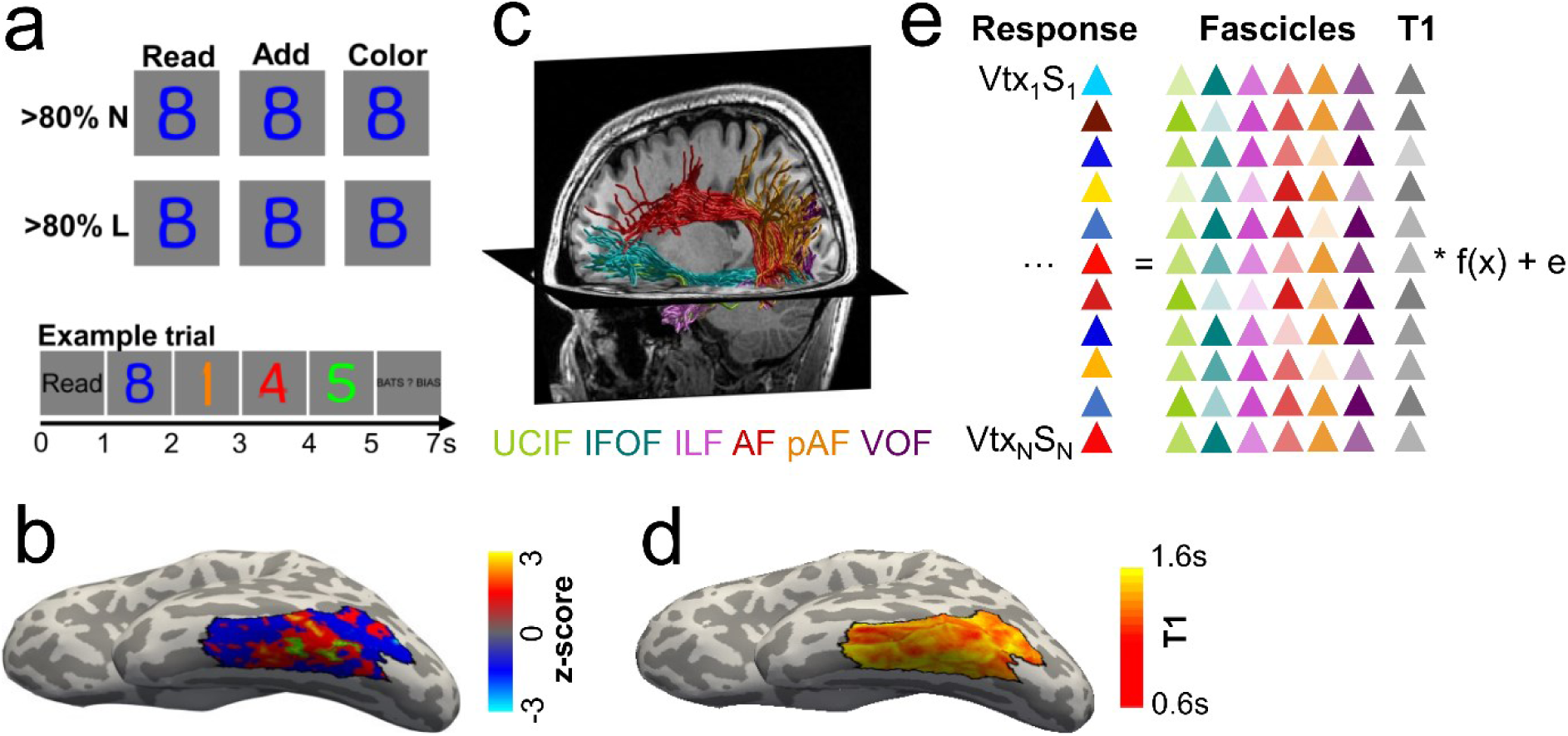
Overview of the experimental approach. **a.** FMRI experiment used to determine reading-related neural responses. Subjects viewed morphs between numbers and letters, containing either >80% letter (<20% number) or >80% number (<20% letter) information. At the beginning of each trial, a cue (Read/Add/Color) indicated which task should be performed, then four stimuli of the same morph type appeared for 1 s each, followed by an answer screen presented for 2 s. Subjects indicated their answer with a button press. Identical stimuli were presented across tasks. Trial structure is shown at the bottom. **b**. Reading-related responses in lateral VTC from fMRI experiment. Data are shown on the inflated cortical surface of a representative subject. We used a t-map (reading> adding + color) to identify a reading-related functional region of interest in the mOTS (green outline, threshold: T≥3), which likely corresponds to the VWFA-2. Finally, the t-map was z-scored to control for inter-individual differences in mean t-values. **c**. White matter fascicles used to predict reading-related responses in lateral VTC. We created a white matter connectome from each subject’s dMRI data and automatically identified six fascicles that have endpoints in the temporal lobe. The endpoint densities of these fascicles were used to predict reading-related responses in lateral VTC. **d**. T_1_ map across lateral VTC estimated from qMRI data shown on the inflated cortical surface of a representative subject. **e**. Schematic of the linear model relating reading-related responses to a weighted sum of white matter fascicle endpoint densities and gray matter T_1_ at each vertex in lateral VTC. Model accuracy was assessed using leave-one-subject-out cross-validation. A*bbreviations:* VTC=ventral temporal cortex, mOTS=mid occipito-temporal sulcus, VWFA=visual word form area, UCIF=uncinate fasciculus, IFOF=inferior frontal occipital fasciculus, ILF=inferior longitudinal fasciculus, AF=arcuate fasciculus, VOF=vertical occipital fasciculus, T1=proton relaxation time, Vtx=vertex, S=Subject.

## 2. Methods

### 2.1 Participants

30 typical adult participants (15 female, mean age ±SE: 27±1 years, 1 left-handed) were recruited from Stanford University and surrounding areas and participated in two experimental sessions. Subjects gave their informed written consent and the Stanford Internal Review Board on Human Subjects Research approved all procedures.

### 2.2 Stimuli and design

In the fMRI experiment, we presented well-controlled character-like stimuli, which could be used for a reading task, a math task, and a color memory task (**Fig. 1a**). These stimuli, which were morphs of numbers and letters, allowed us to map reading-related responses while keeping the visual input constant (Grotheer et al., 2019, 2018). At the beginning of each trial, subjects were presented with a cue (Add, Read, or Color), indicating which task they should perform. In the reading task, subjects were instructed to read the word in their head, and to indicate which word had been presented. In the adding task, participants were asked to sum the values of the stimuli and to indicate the correct sum. Finally, in the color task, participants were asked to memorize the color of the stimuli and to indicate which color was shown during the trial. After the cue, 4 images were shown sequentially, followed by an answer screen. All images in a trial were either number morphs (N, >80% number + <20% letter) or letter morphs (L, >80% letter + <20% number), i.e. stimuli that mostly contained information from one category, but held just enough evidence from the other category to be recognizable as both letters and numbers. The same stimuli appeared in all tasks. The answer screen was presented for 2 seconds and showed the correct answer as well as one incorrect answer at counterbalanced locations left and right of fixation. Participants performed 6 runs, each lasting six minutes, and the task order was randomized across runs and participants. Prior to the experiment, subjects were given training to ensure that they could perform the task with at least 80% accuracy.

### 2.3 Functional MRI data acquisition and preprocessing

fMRI data was collected at the Center for Cognitive and Neurobiological Imaging at Stanford University, using a GE 3 tesla Signa Scanner with a 32-channel head coil. We acquired 48 slices covering the entire cortex using a T2*-sensitive gradient echo sequence (resolution: 2.4 mm × 2.4 mm × 2.4 mm, TR: 1000 ms, TE: 30 ms, FoV: 192 mm, flip angle: 62°, multiplexing factor of 3). A subset (N=20) of the fMRI data were used for previous studies (Grotheer et al., 2019, 2018).

A whole-brain, anatomical volume was also acquired, once for each participant, using a T1-weighted BRAVO pulse sequence (resolution: 1mm × 1 mm × 1 mm, TI=450 ms, flip angle: 12°, 1 NEX, FoV: 240 mm). The anatomical volume was segmented into gray and white matter using FreeSurfer (http://surfer.nmr.mgh.harvard.edu/), with manual corrections using ITKGray (http://web.stanford.edu/group/vista/cgi-bin/wiki/index.php/ItkGray). From this segmentation, each participant’s cortical surface was reconstructed. Each participant’s anatomical brain volume was used as the common reference space for all analyses, which were always performed in individual native space. In order to be able to derive linear models specific to lateral VTC, we first drew an anatomical boundary of lateral VTC in the FreeSurfer average brain. This anatomical boundary stretched from the mid-fusiform sulcus (medial boundary) to the inferior temporal gyrus (lateral boundary) and from the anterior end of the occipito-temporal sulcus (OTS, anterior boundary) to the posterior transverse collateral sulcus (posterior boundary). This boundary was chosen to match previous work (Nordt et al., 2019) except for the anterior boundary, which we shifter more anteriorly to ensure that the entire OTS is included. This anatomical boundary, which is made available in GitHub (https://github.com/VPNL/predictFuncFromStructCode), was transformed to each individual’s native space (**black outline in Fig. 1b**,**c**) and all further analyses were restricted to this boundary.

The functional data was analyzed using the mrVista toolbox (http://github.com/vistalab) for Matlab, as in previous work (Grotheer et al., 2019, 2018). The data was motion-corrected within and between scans and then manually aligned to the anatomical volume. The manual alignment was optimized using robust multiresolution alignment (Nestares and Heeger, 2000). No smoothing was applied. The time course of each voxel was high-pass filtered with a 1/20 Hz cutoff and converted to percentage signal change. A design matrix of the experimental conditions was created and convolved with the hemodynamic response function (HRF) implemented in SPM (http://www.fil.ion.ucl.ac.uk/spm) to generate predictors for each experimental condition. Response coefficients (betas) were estimated for each voxel and each predictor using a general linear model (GLM).

### 2.3 Functional regions of interest and functional activation maps

We evaluated reading-related responses by contrasting the reading task with the adding and the color task in the experiment. The resulting T-maps were used to identify a reading-related functional region of interest (fROI) in the left mid occipitotemporal sulcus (mOTS) (**green outline in Fig. 1b**). This region, which likely corresponds to the anterior segment of the visual word form area (Cohen et al., 2000; Dehaene and Cohen, 2011) or VWFA-2 (Grotheer et al., 2018; Lerma-Usabiaga et al., 2018; Stigliani et al., 2015; Weiner et al., 2017), was defined in each participant’s cortical surface using both functional and anatomical criteria. Specifically, we took only those vertices that (i) showed a preference for the reading task over the other tasks beyond the threshold of T≥3 and (ii) fell within the mOTS. The resulting VFWA-2 fROI could be identified in 28/30 participants (fROI size±SE: 301±52 mm^3^).

Following this, each subject’s reading-related T-map was z-scored to control for inter-subject variability in overall response magnitude and mapped onto that individual’s inflated cortical surface using FreeSurfer (**Fig. 1b**). Responses at each vertex within the anatomical boundary of lateral VTC were used to derive linear models to predict reading-related responses based on the brain’s white matter fascicles and gray matter tissue microstructure. As activations during reading were lower and less frequent in the right than the left hemisphere, which is consistent with the current literature (Behrmann and Plaut, 2015; Grotheer et al., 2019), here we focus on the left hemisphere only.

### 2.5 Diffusion MRI data acquisition and processing

Diffusion-weighted MRI (dMRI) data was collected from the same participants during a different day than the fMRI data, at the same facility and with the same 32-channel head-coil. DMRI was acquired using a dual-spin echo sequence in 96 different directions, 8 non-diffusion-weighted (b=0) images were collected, 60 slices provided full head coverage (resolution: 2 mm × 2 mm × 2 mm, TR: 8000 ms, TE: 93.6 ms, FoV: 220 mm, flip angle: 90°, b: 2000 s mm^−2^).

DMRI data was preprocessed using a combination of tools from mrTrix3 (github.com/MRtrix3/mrtrix3) and mrDiffusion toolbox (http://github.com/vistalab/vistasoft) for Matlab. First, we denoised the data using i) a principal component analysis, ii) Rician based denoising, and iii) Gibbs ringing corrections (Kellner et al., 2016; Veraart et al., 2016b, 2016a). Second, we corrected for eddy currents and motion using FSL (Smith et al., 2004) (https://fsl.fmrib.ox.ac.uk/), and we performed bias correction using ANTs (Tustison et al., 2010). Third, dMRI data was registered to the average of the non-diffusion weighted images and aligned to the corresponding high-resolution anatomical brain volume using rigid body transformation. Fourth, voxel-wise fiber orientation distributions (FODs) were calculated using constrained spherical deconvolution (CSD) (Tournier et al., 2007). For this, we used the Dhollander algorithm to estimate the three-tissue response function (Dhollander et al., 2016) with automated selection of the number of spherical harmonics. The FODs were then used for tractography.

We used MRtrix3 (Tournier et al., 2019) (RC3, http://www.mrtrix.org/) to generate a white matter connectome for each subject. For each connectome, we used probabilistic fiber tracking with the following parameters: algorithm: IFOD1, step size: 0.2 mm, minimum length: 4 mm, maximum length: 200 mm, FOD amplitude stopping criterion: 0.1, maximum angle: 13.5°. We used anatomically constrained tractography (ACT) (Smith et al., 2012), which utilizes information of different tissue types from the FreeSurfer (https://surfer.nmr.mgh.harvard.edu/) segmentation of each participant’s high-resolution anatomical scan to optimize tractography. ACT also allowed us to identify the gray-white matter interface (GWMI). Seeds for tractography were randomly placed within this interface within the anatomical boundary of lateral VTC. This enabled us to ensure that we based our predictions on tracts that reach the gray matter. Each connectome consisted of 3 million streamlines.

Finally, we used Automated Fiber Quantification (AFQ, Yeatman et al., 2012, https://github.com/yeatmanlab/AFQ) to automatically segment the connectome of each participant into well-established fascicles. The resulting classified connectome was optimized by removing tracts that were located more than 4 standard deviations away from the mean of their respective fascicle (similar to Yeatman et al., 2014, 2012). We conducted all subsequent analyses on these classified white matter tracts, as we were interested in identifying which fascicles predict the spatial layout of reading-related responses in lateral VTC. We focused on those 6 fascicles that connect the temporal lobe with other parts of the brain and therefore may originate or end in lateral VTC (**Fig. 1c**). These tracts are: i) the uncinate fasciculus (UCIF), ii) the inferior frontal occipital fasciculus (IFOF), iii) the inferior longitudinal fasciculus (ILF), iv) the arcuate fasciculus (AF), v) the posterior arcuate fasciculus (pAF) and vi) the vertical occipital fasciculus (VOF). In order to determine which of these fascicles are predictive of reading-related responses, we first mapped the endpoint density of each fascicle onto the cortical surface and then used the endpoint densities at each vertex as predictors in linear models of reading-related responses.

### 2.6 Quantitative MRI data acquisition and preprocessing

Quantitative MRI (qMRI, Mezer et al., 2013) data was collected within the same session and with the same head coil as the dMRI data. T_1_ relaxation times were measured from four spoiled gradient echo images with flip angles of 4°, 10°, 20° and 30° (TR: 14 ms, TE: 2.4 ms). The resolution of these images was later resampled from 0.8×0.8×1.0 mm^3^ to 1mm isotropic voxels, and qMRI data was aligned with the high-resolution anatomical scan using rigid body transformation. We also collected four additional spin echo inversion recovery (SEIR) scans with an echo planar imaging read-out, a slab inversion pulse and spectral spatial fat suppression (TR: 3 s, resolution: 2 mm × 2 mm × 4 mm, 4 echo time set to minimum full, 2x acceleration, inversion times: 50, 400, 1200, and 2400 ms). The purpose of these SEIRs was to remove field inhomogeneities.

Both the spoiled gradient echo and the SEIR scans were processed using the mrQ software package (https://github.com/mezera/mrQ) for Matlab to estimate the proton relaxation time (T_1_) in each voxel, as in previous studies (Gomez et al., 2017; Grotheer et al., 2019; Mezer et al., 2013; Yeatman et al., 2014a). The mrQ analysis pipeline corrects for RF coil bias using the SEIRs scans, which produces accurate proton density (PD) and T_1_ fits across the brain. The T_1_ map of each subject was co-registered to the same anatomical whole-brain volume as dMRI and fMRI data and mapped to the inflated cortical surface of each participant (**Fig. 1d**). We used the T_1_ maps to evaluate if gray matter tissue microstructure contributes to the prediction of reading-related responses. For this, T_1_ at each vertex within the anatomical boundary of lateral VTC was used as an additional predictor in linear models of reading-related responses.

### 2.7 Linear model: feature selection

In order to determine if white and gray matter anatomy predicts the spatial layout of reading-related responses, we derived linear models that relate the endpoint density of fascicles as well as gray matter T_1_ to reading-related responses. First, using a randomly selected subset of 10 out of 30 participants (the feature selection set), we evaluated which white and gray matter features best predict reading-related responses in lateral VTC. During this feature selection, we evaluated which white matter fascicles are predictive of the map of reading-related responses by deriving separate linear models relating the reading-related response map to each fascicle’s endpoints in lateral VTC. We tested how well the endpoint density of each fascicle at each vertex within the anatomically defined lateral VTC predicts reading-related responses using leave-one-subject-out cross-validation. That is, we derived the model using data of all but one subject and then predicted responses in the left-out subject, iterating across all participants. In order to assess the performance of the model, we then measured the correlation between the predicted to the measured reading-related response map in the left-out subject. We used Bonferroni corrected paired t-tests to evaluate if the correlation between the predicted and measured reading-related responses is significantly greater than 0. Finally, those fascicles that lead to significant predictions were combined into a new linear model that we will refer to as the combined fascicle model.

We tested if different fascicles capture independent variance in the reading-related responses by evaluating if the combined fascicle model (a weighted sum of the endpoints of the AF, ILF and VOF) outperforms the best individual fascicle model. For this, first, we compared predicted and measured reading-related responses in leave-one-subject-out cross-validations, as described above. We used Bonferroni corrected paired t-tests to evaluate if the correlation between the predicted and measured reading-related responses is significantly greater than 0 in each model. Next, we derived both models using the data of all 10 participants and formally tested if the combined fascicle model outperforms the best individual fascicle model using a simulated likelihood ratio test with 1000 simulations. Given that this analysis indicated the combined fascicle model to outperform the individual fascicle models, we used the combined fascicle model in further steps.

After determining which fascicles are predictive of reading-related responses, in the last step of the feature selection, we assessed whether information about tissue properties in the gray matter, i.e., T_1_ at each vertex, improves the prediction of reading-related responses relative to the combined fascicles model. Thus, we added T_1_ as an additional predictor to the combined fascicle model and compared predicted and measured reading-related responses in leave-one-subject-out cross-validations for the combined fascicle model and the combined fascicle+T_1_ model. We used Bonferroni corrected paired t-tests to evaluate if the correlation between the measured reading-related responses and those predicted by each model is significantly greater than 0. We tested if the model that includes T_1_ outperforms the combined fascicle model using a simulated likelihood ratio test with 1000 simulations. Given that these analyses indicated T_1_ at each vertex to be an informative predictor, we added this predictor to our final model, which we refer to as fascicles+T_1_.

### 2.8 Linear model: model validation

To ensure independence of feature selection and model testing, we tested the predictiveness of the fascicles+T_1_ model in the remaining 20 participants (validation set). For this, at each vertex, we compared the measured reading-related responses with the responses predicted by the fascicles+T_1_ model using leave-one-subject-out cross-validation in the independent subjects. We used a paired t-test to evaluate if the correlation between the predicted and measured reading-related responses is significantly greater than 0. We also assessed whether the predictiveness of our model depends on the selectivity of each vertex. Thus, in each participant’s lateral VTC we identified the 10, 20 and 30% vertices with the strongest preference for reading, those with the strongest preference against reading and those with no strong preference. These sets of vertices where identified in both the measured and the predicted reading-related response maps. Then, we assessed the spatial overlap between measured and predicted vertices using the dice coefficient DC (Dice, 1945): 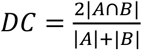, where |A| is the measured vertices, |B| is the predicted vertices and |A∩B| is the intersection between these two sets of vertices. Additionally, we calculated what is the chance level DC by randomly selecting 10, 20 and 30% of lateral VTC vertices in the predicted map and calculating the overlap between these randomly chosen vertices and those vertices that show the strongest preference for reading in the measured reading-related responses. Finally, we tested the significance of the prediction by comparing the measured DCs to the chance level DCs for the 20% sets of vertices, using Bonferroni corrected paired t-tests.

### 2.9 Linear Discrimination Analysis

We also tested how well the anatomical features chosen above predict the location of the VWFA-2 in the validation set. For this, we used a linear discrimination analysis (LDA) and derived a linear classifier that uses fascicle endpoints and T_1_ to determine which vertices in lateral VTC are within the VWFA-2 and which are outside this fROI. We used a leave-one-subject-out cross-validation approach iterating across all 20 subjects. To determine the accuracy of this classifier, first, we visually compared the predicted and the measured VWFA-2 fROIs on the cortical surface of each held-out subject. Next, we assessed the prediction accuracy by quantifying the spatial overlap between the predicted and the measured VWFA-2 using the DC. We also performed a similar analysis using a 7mm disk ROI (radius was chosen to match the average radius of the fROIs) placed in the center of the VWFA-2 to test whether out model predicts only the location of the VWFA-2 or also its shape. We reasoned that, if white and gray matter anatomy also predict the shape of the VWFA-2, the prediction of the disk ROI should be significantly worse than the prediction of the VWFA-2 fROI. Finally, we generated a chance level DC by randomizing the vertices that are within and outside of the predicted VWFA-2 and quantifying the spatial overlap between these randomly chosen vertices and the measured VWFA-2.

Finally, to assess if the predicted VFWA-2 responds more strongly to the reading task than the adding and color tasks, i.e. shows the desired preference for reading, we extracted the responses of the predicted VWFA-2 during the fMRI experiment in each held-out subject. We used repeated measures ANOVA’s with task and stimulus as factors to evaluate the effect of the performed task and the stimulus on neural responses within the predicted VWFA-2 fROI.

### 2.10 Data and code availability

The fMRI and qMRI data were analyzed using the open source mrVista software (available in GitHub: http://github.com/vistalab/vistasoft) and mrQ software (available in GitHub: https://github.com/mezera/mrQ) packages, respectively. The dMRI data were analyzed using open source software, including MRtrix3 (Tournier et al., 2012) (http://www.mrtrix.org/) and AFQ (Yeatman et al., 2012b) (https://github.com/yeatmanlab/AFQ). We make the entire pipeline freely available; custom code for preprocessing, tractography and further analyses are available in github (https://github.com/VPNL/fat). Code for reproducing all figures and statistics are made available in github as well (https://github.com/VPNL/predictFuncFromStructCode). The data generated in this study will be made available by the corresponding author upon reasonable request.

## 3. Results

### 3.1 Endpoints of the ILF, AF, and VOF predict reading-related responses

In the current study we tested the hypothesis that the white matter fascicles of the human brain and the cortical microstructure (T_1_) predict where reading-related responses fall in lateral ventral temporal cortex (lateral VTC) in a given adult individual. Thus, we related white and gray matter anatomy to reading-related responses in two steps: 1) We identified the anatomical features that best predict reading-related responses (feature selection), and 2) we developed a linear model based on the most informative features and tested how well this model predicts the map of reading-related responses in lateral VTC as well as the location of the VFWA in new participants (model validation). In order to ensure independence of feature selection and model validation, we split our data into two groups of participants: 10 participants were used for feature selection and 20 other participants were used for model validation.

During feature selection, we first evaluated which white matter fascicles predict reading-related responses in lateral VTC. To do so, we i) mapped reading-related responses across lateral VTC in each individual by contrasting responses elicited by a reading task with those elicited by color and math tasks performed on identical visual stimuli (Grotheer et al., 2019, 2018), ii) mapped the endpoint density of those fascicles that connect to the temporal lobe across lateral VTC, iii) quantitatively evaluated if there is a relationship between reading-related responses and the endpoints of each of the fascicles using linear models that relate reading-related response magnitude to fascicle endpoint density, and (iv) tested if a linear model which combines the endpoint densities of multiple fascicles better explains reading-related responses than a model based on individual fascicles.

Six major fascicles connect to the temporal lobe: i) the uncinate fasciculus (UCIF), ii) the inferior frontal occipital fasciculus (IFOF), iii) the inferior longitudinal fasciculus (ILF), iv) the arcuate fasciculus (AF), v) the posterior arcuate fasciculus (pAF), and vi) the vertical occipital fasciculus (VOF). We automatically identified these fascicles in each individual and mapped their endpoint density to the cortical surface within an anatomical boundary of lateral VTC. We then visually inspected the resulting endpoint density maps (**Fig. 2a** shows each of these maps in a representative subject). Across subjects, we found that i) the endpoints of the ILF, AF, and pAF are distributed across most of lateral VTC, ii) the VOF and IFOF have endpoints only in the posterior end of lateral VTC, and iii) the UCIF rarely has any endpoints in lateral VTC.

**Fig. 2.**
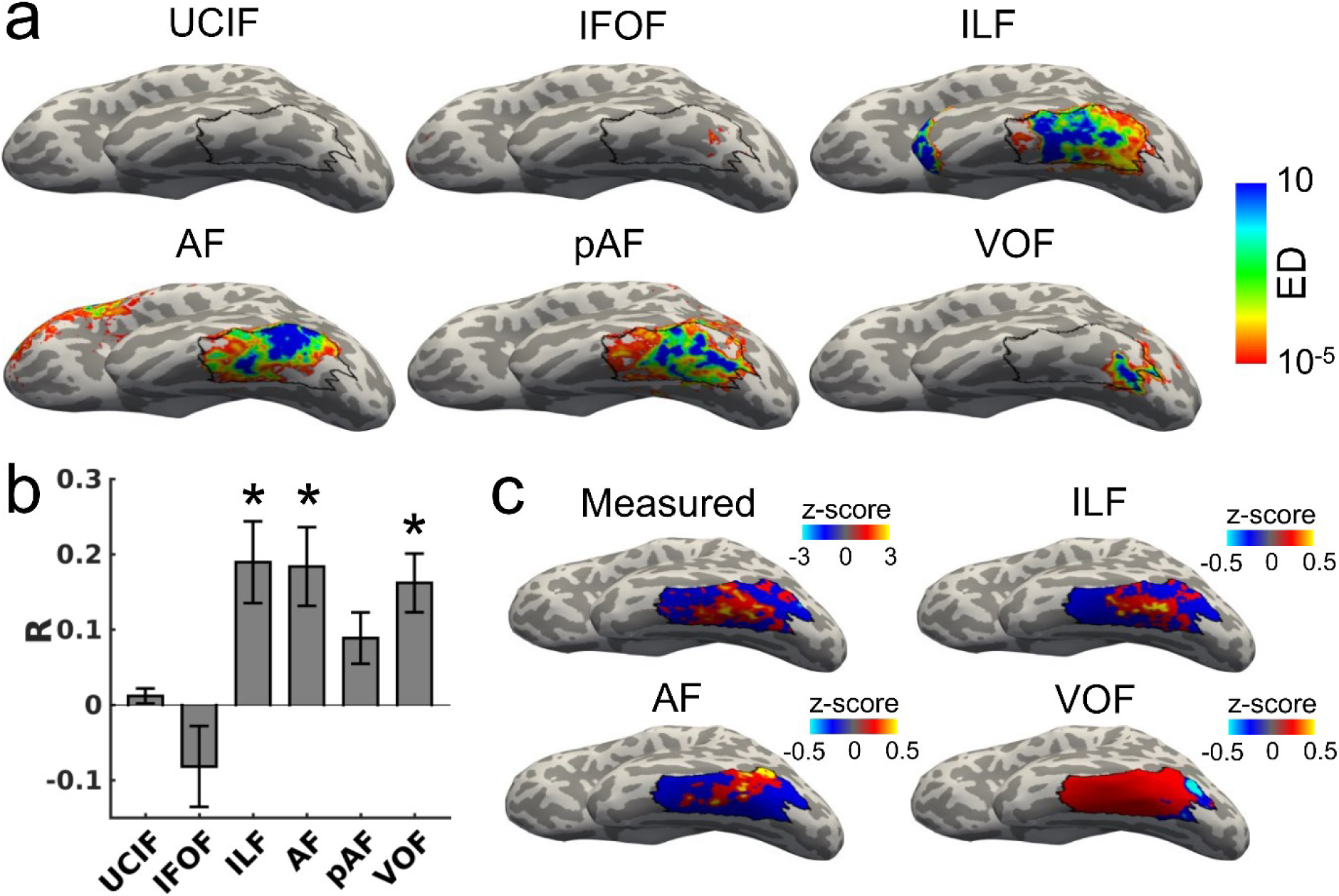
Endpoints of ILF, AF, and VOF predict reading-related responses in lateral VTC. **a.** Maps of endpoint density for UCIF, IFOF, ILF, AF, pAF, and VOF, presented on the inflated cortical surface of a representative participant. **b**. The endpoint density of each fascicle was used to train a linear model to predict reading-related responses. Figure shows to what degree predicted and measured reading-related responses correlated in leave-one-subject-out cross-validation. *=significant correlation between predicted and measured reading-related responses in left-out subject (R significantly above 0; p≤0.008, Bonferroni corrected). **c**. Maps of predicted reading-related responses for each of the fascicles that showed significant predictions, presented on the inflated cortical surface of a representative subject. Abbreviations: UCIF=uncinate fasciculus, IFOF=inferior frontal occipital fasciculus, ILF=inferior longitudinal fasciculus, AF=arcuate fasciculus, pAF=posterior arcuate fasciculus, VOF=vertical occipital fasciculus.

To determine which of these fascicles are predictive of reading-related responses, we next derived linear models that relate reading responses in lateral VTC to endpoint density separately for each of these fascicles. We trained one model for each fascicle and evaluated the performance of the model using leave-one-subject-out cross-validation. That is, we trained the model on 9/10 participants and then tested how well the model predicts reading-related responses in the held-out participant, iterating across all leave-one-out combinations. We evaluated each model’s performance by correlating the predicted responses with the measured reading-related responses in each held-out participant.

Results of this analysis show that the endpoint density of the ILF, AF, and VOF predict reading-related responses in lateral VTC. That is, linear models relating the endpoints of each of these fascicles to reading-related responses show a significant cross-validated correlation between the predicted and measured responses (paired two-sided t-tests against 0, with a Bonferroni adjusted threshold of p≤0.008; **Fig. 2b**; mean R±SE: ILF: 0.19±0.05, AF: 0.18±0.05, VOF: 0.16±0.04).

To better understand the relationship between each fascicle’s endpoint density and reading-related responses, next, we compared the endpoint density maps described above (**Fig. 2a** shows these maps in a representative subject) with the predicted reading-related responses (**Fig. 2c** shows a representative subject) for each fascicle that showed significant predictions. We find that regions in lateral VTC that have a high endpoint density for the AF and ILF (blue in **Fig. 2a**) coincide with regions that show a preference for reading (red in **Fig. 2c**-measured). In contrast, regions that show high endpoint density for the VOF, which predominately showed endpoints in the posterior end of lateral VTC, showed a negative preference for reading.

Our data suggests that the endpoints of the ILF, AF, and VOF can predict reading-related responses in lateral VTC. However, an open question is whether these fascicles provide overlapping or complimentary predictions of reading-related responses. The latter seems plausible as the endpoint densities of these three fascicles have different distributions across lateral VTC (**Fig. 2a**) and generated different predictions of reading-related responses (**Fig. 2c**).

To test these possibilities, we generated a new linear model that relates responses during reading to a weighted combination of ILF, AF, and VOF endpoints in lateral VTC. Then, we compared the performance of this combined model and the best individual fascicle model using leave-one-subject-out cross-validation in the 10 subjects used for feature selection. We find that a linear model with three fascicles improved prediction of reading-related responses. That is, there is a higher correlation between the predicted and measured reading-related responses compared to a linear model based on a single fascicle (mean R±SE: best individual fascicle (ILF): 0.19±0.05, combined fascicle model: 0.27±0.03; **Fig. 3a**; example prediction in a representative individual in **Fig. 3b**). This improvement is statistically significant (p=0.001, simulated likelihood ratio test comparing models derived from the data of all 10 participants).

**Fig. 3.**
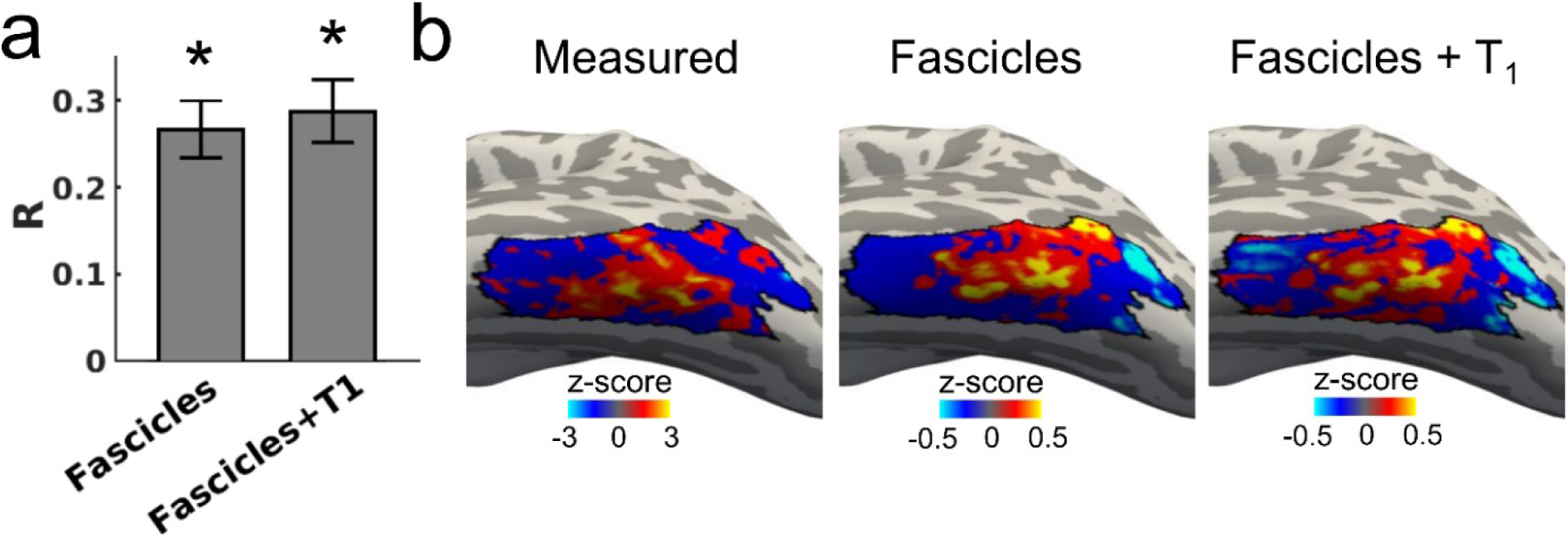
Linear model based on fascicle endpoint densities and cortical T_1_ best predicts reading-related responses in lateral VTC. **a.** Correlation between predicted and measured reading-related responses for linear models that combine the endpoint densities of the ILF, AF, and VOF (the combined fascicle model) or the endpoint densities of the ILF, AF, and VOF as well as cortical T_1_ (the fascicle + T_1_ model). *=Significant correlation between predicted and measured reading-related responses in leave-one-subject-out cross-validation (R significantly above 0; p≤0.025, Bonferroni corrected). Model comparison showed that the fascicle + T_1_ model outperforms the combined fascicle model (likelihood ratio test, p=0.001). **b**. Maps of measured and predicted reading-related responses for each model, presented on the inflated cortical surface of a representative subject. Abbreviations: T_1_=proton relaxation time, ILF=inferior longitudinal fasciculus, AF=arcuate fasciculus, VOF=vertical occipital fasciculus.

Overall, this analysis indicates a consistent spatial relationship between the endpoint densities of the AF, ILF, and VOF and reading-related responses in lateral VTC.

### 3.2 Cortical tissue properties improve prediction of reading-related responses

Next, as the last step of feature selection, we tested whether cortical tissue microstructure in lateral VTC, assessed by T_1_ relaxation time, further improves the prediction of reading-related responses compared to the combined fascicle model. We reasoned that gray matter microstructure may covary with reading-related responses and that regions with high reading-related responses may show a different microstructure than other parts of lateral VTC.

First, using qMRI, we estimated T_1_ at each vertex in lateral VTC. Next, we added the T_1_ values as an additional predictor to the combined fascicle model. Using the same 10 subjects as in the prior analyses, we trained the fascicle+T_1_ model on 9/10 participants and then tested how well it predicts reading-related responses in the held-out subject, iterating across all leave-one-out combinations. We found that this model’s prediction of reading-related responses was significantly correlated with the measured responses in the left out subjects (**Fig. 3a**, paired two-sided t-tests with a Bonferroni adjusted threshold of p≤0.025), which is also evident when visually comparing the spatial layout of the measured reading-related responses with the predicted responses (**Fig. 3b** shows a representative subject). It should be noted though, that the fascicles that are included in the fascicle+T_1_ model were chosen based on the same 10 subjects that the model was then tested on, we will further investigate the fascicle+T_1_ model’s performance on independent data in the next sections.

Crucially, we found that adding gray matter T_1_ to the combined fascicle model improved the prediction of reading-related responses in lateral VTC (mean R±SE: combined fascicle model: 0.27±0.03, combined fascicle + T_1_ model: 0.29±0.04). This improvement was statistically significant (p=0.001, simulated likelihood ratio test comparing models derived from the data of all 10 participants). Results of this analysis suggests that cortical T_1_ explains independent variance in the spatial layout of reading-related responses, which is not captured by the white matter fascicles alone.

Together, these analyses suggest that the endpoint density of the AF, ILF and VOF as well as cortical microstructure assessed with T_1_ show a consistent spatial relationship with reading-related responses in lateral VTC across individuals.

### 3.3 Model based on fascicle endpoints and cortical T_1_ predicts both the peaks and the troughs of reading-related responses

To further assess the robustness of the fascicles+T_1_ model, during model validation, we tested how well this model predicts reading-related responses in 20 new participants, particularly focusing on how well the model predicts i) the spatial layout of peaks and troughs of reading-related responses as well as ii) the location of the visual word form area, a region that is considered critical for reading and word recognition.

First, to test how well the combined model – the linear model that uses AF, ILF, VOF and cortical T_1_ as predictors – captures the map of reading-related responses in independent data, we again used leave-one-subject-out cross-validation. That is, we trained the combined model on 19/20 new participants and then tested how well it predicts reading-related responses in the held-out subject, iterating across all leave-one-out combinations. Results reveal that, on average, the predicted and the measured reading-related responses in each held-out subject correlate significantly above chance (**Fig. 4a shows measured and predicted maps side-by-side in 3 example subjects**; mean R±SE: 0.28±0.03; mean R was significantly greater than 0, paired t-test, p<0.0001) confirming the predictive nature of these combined features of the white and gray matter in 20 new participants.

**Fig. 4.**
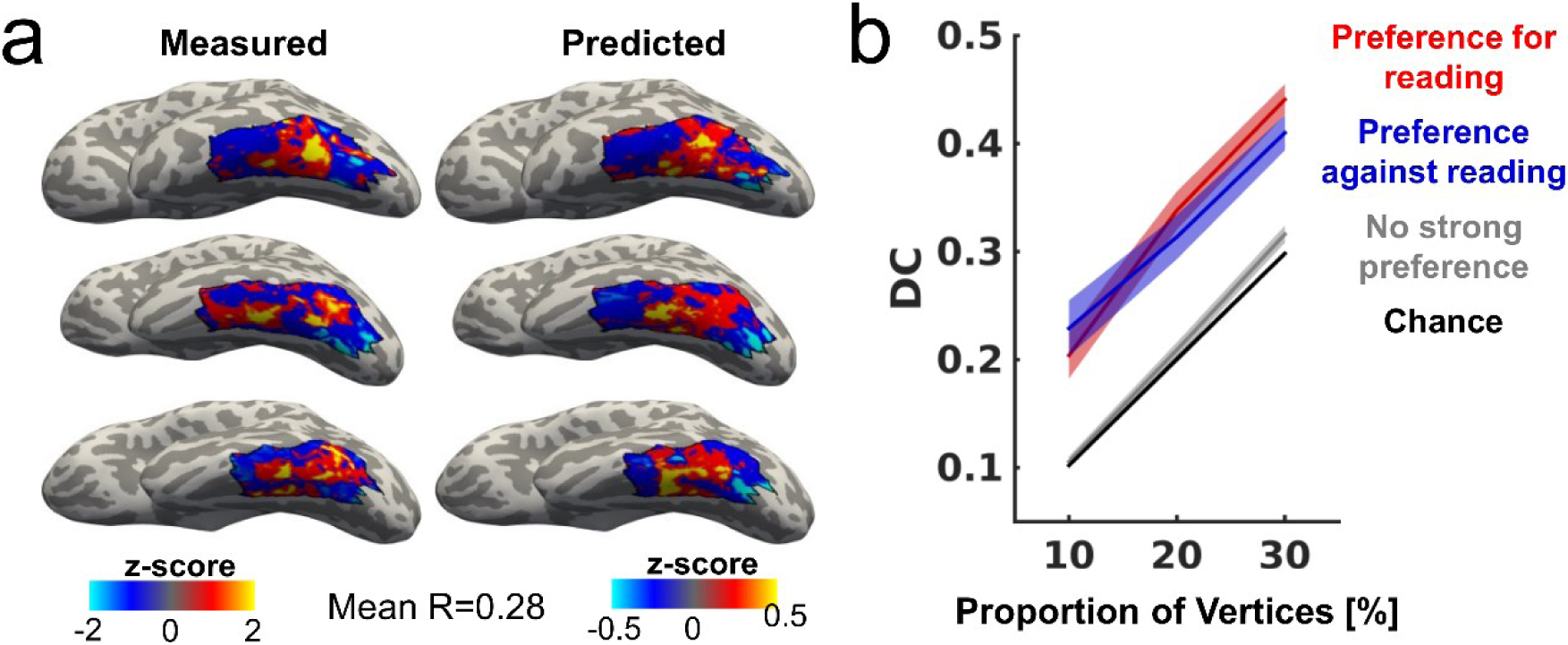
Endpoints of AF, ILF, and VOF in combination with cortical T_1_ predict high and low reading-related responses in lateral VTC. **a**. Comparison of measured and predicted maps of reading-related responses in three example participants. The predicted maps were created using leave-one-subject-out cross-validation with a linear model that combines the endpoint density of the ILF, AF, and VOF with cortical T_1_. **b**. DC analysis comparing the predicted and the measured maps. Analysis shows that the linear model significantly predicts the location of both the peaks and the troughs of reading-related responses but fails to predict vertices with no strong preference (p<0.01, Bonferroni corrected). *Abbreviations:* T_1_=proton relaxation time, ILF=inferior longitudinal fasciculus, AF=arcuate fasciculus, VOF=vertical occipital fasciculus, DC=dice coefficient.

Next, we assessed whether the fascicles+T_1_ model best predicts the vertices with the strongest reading-related responses (referred to as vertices with a preference for reading), or whether it predicts vertices that have a preference for the other tasks (preference against reading) or those with no strong preference equally well. Thus, in each participant’s lateral VTC, we identified the top 10%, 20% and 30% vertices that show the strongest preference for reading, both for the measured reading-related responses and for the reading-related responses predicted by the fascicles+T_1_ model. Then, we quantified their overlap using the dice coefficient (DC, Dice, 1945). We found substantial spatial overlap between the measured and the predicted 10, 20, and 30% vertices with the strongest preference for reading (**Fig. 4b-red**, DC±SE: 10%: 0.20±0.02, 20%: 0.34±0.02, 30%: 0.44±0.01; paired two-sided t-tests performed on 20% above chance with a Bonferroni adjusted threshold of p≤0.01), suggesting that this model accurately captures those vertices that show a strong preference for reading, or in other words, the peaks of reading-related responses.

Next, we evaluated if the significant overlap between measured and predicted responses is specific to those vertices with the highest reading-related responses or if the fascicles+T_1_ model also predicts those vertices with the lowest reading-related responses, i.e. vertices that show higher responses during the adding and color tasks than during the reading task. We followed the procedure as described above and found high spatial overlap between the measured and the predicted 10%, 20% and 30% vertices with a preference against reading (**Fig. 4b-blue**, DC±SE: 10%: 0.23±0.03, 20%: 0.31±0.02, 30%: 0.41±0.02; paired two-sided t-tests performed on 20% above chance with a Bonferroni adjusted threshold of p≤0.01), suggesting that the fascicles+T_1_ model accurately predicts the troughs of reading-related responses.

Finally, we also tested how well the model predicts vertices that have neither a strong preference for reading or a strong preference against reading. In contrast to vertices with strong positive or negative preferences, the prediction of 10, 20, 30% vertices with weak preferences did not differ from chance (**Fig. 4b-gray**, DC±SE: 10%: 0.11±0.005, 20%: 0.21±0.007, 30%: 0.32±0.008; paired two-sided t-tests performed on 20% not significantly different from chance). Together, these data show that our model accurately captures both the peaks and the troughs of reading-related responses in lateral VTC.

### 3.4 Fascicle endpoints and cortical T_1_ predict the location of the visual word form area

The visual word form area (VWFA, Cohen et al., 2000; Dehaene and Cohen, 2011), a functional region of interest (fROI) in the occipito-temporal cortex (OTS), plays a critical role in reading (Gaillard et al., 2006). The VFWA has two clusters in the left hemisphere: one on the posterior OTS (also referred to as pOTS-words/VWFA1) and one on the mid OTS (also referred to as mOTS-words/VWFA-2) (Lerma-Usabiaga et al., 2018). Since our reading-related ROI coincides with VWFA-2 (Grotheer et al., 2018), we tested if our fascicles+T_1_ model can also accurately predict the location and boundaries of this region in individual subjects.

To this end, first we identified the VWFA-2 by selecting those voxels in the mOTS that show significantly higher responses in the reading task than the adding and the color tasks (threshold: T≥3, voxel-level) in each of the 20 participants used for model testing. We then performed linear discrimination analyses (LDAs) with a combined fascicles+T_1_ classifier to distinguish vertices that are part of the VWFA-2 from those that are not part of the VWFA-2. As before, we used 20-fold leave-one-subject-out cross-validation. That is, we trained the classifier on 19 participants and predicted the fROI in the held-out subject, iterating across all possible leave-one-out combinations. We then assessed the accuracy of the classifier by i) visually comparing the predicted fROI with the measured fROI in each held-out subject, ii) quantifying the spatial overlap between the measured and predicted fROI using the DC, and iii) evaluating the response characteristics of the predicted fROI to test if it shows the expected preference for the reading task compared to the color and adding tasks in the experiment.

Within each participant, we first plotted the measured and the predicted VWFA-2 side-by-side and visually inspected their spatial correspondence. We find that there is varying overlap between the predicted and the measured VWFA-2 across subjects, with some participants showing almost perfect overlap and some subjects showing more misclassifications (**Fig. 5a** visualizes this range by showing the participants with the most overlap, the least overlap and average overlap).

**Fig. 5.**
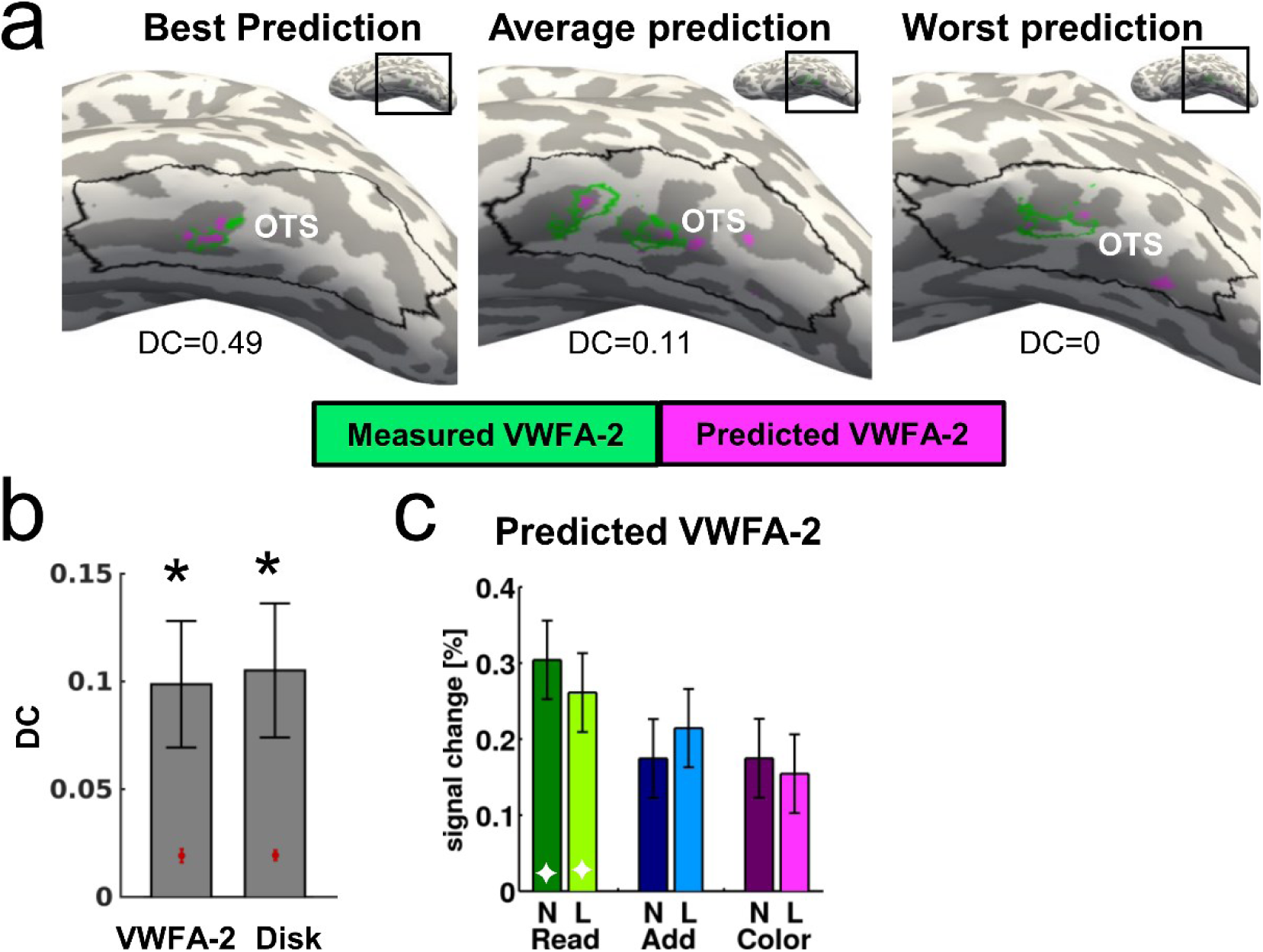
Endpoints of AF, ILF, and VOF in combination with cortical T_1_ predict the location the VWFA-2. **a**. Comparison of predicted and measured VWFA-2 in three representative participants. The fROI was predicted using LDA with leave-one-subject-out cross-validation and a model that combines T_1_ with the endpoint densities of the ILF, AF and VOF. **b**. Dice coefficient (DC) analysis quantifying the spatial overlap between predicted and the measured fROIs for i) the VWFA-2 (left) and ii) a 7mm disk placed in the center of the VWFA-2 (right). *= DC significantly greater than chance, p≤0.03, Bonferroni corrected; chance level indicated by red circle. **c**. Mean responses of the predicted VWFA-2 during the reading, adding and color tasks performed on number morphs (N) and letter morphs (L). ◊= significant main effect of task, repeated measures ANOVA, p≤0.05). *Abbreviations:* fROI=functional region of interest, LDA= linear discrimination analysis, T_1_=proton relaxation time, ILF=inferior longitudinal fasciculus, AF=arcuate fasciculus, VOF=vertical occipital fasciculus, DC=dice coefficient, mOTS=mid occipito-temporal sulcus.

Interestingly, even in participants with average or no overlap between measured and predicted fROI, the predicted fROI still generally falls within the anatomical boundaries of the mOTS and hence in a biologically plausible location.

Next, we used the DC to quantify the spatial overlap between the predicted and the measured VWFA-2 in each participant and compared the measured DC to the chance level DC. Results show that the fascicles+T_1_ model predicts the location of the VWFA-2 significantly above chance (paired t-test, p=0.01, significantly above Bonferroni corrected threshold of p<0.03, **Fig. 5b**). In order to evaluate if the combined model is informative only about the location or also about the shape of the fROI, we also performed the same analysis with a 7mm disk ROI placed in the center of each participant’s VWFA-2 fROI. We found that the overlap between the predicted and the measured disk ROI is significantly above chance (paired t-test, p=0.01, significantly below Bonferroni corrected threshold of p<0.03, **Fig. 5b**) and that there is no difference in prediction accuracy between the VWFA-2 and the disk fROI (paired t-test, p=0.61). This suggests that the combined model accurately predicts the location, but not the shape of the reading-related fROI in mOTS.

Finally, we tested if the predicted VWFA-2 shows the expected preference for the reading task over the adding and color tasks. To this end, we extracted the responses for each of the tasks from the predicted fROI in each participant. Results show that responses in the predicted VWFA-2 were higher during the reading than the color and the adding tasks (**Fig 5c**). This difference was statistically significant (main effect of task, repeated measures ANOVA with task and stimulus as factors: F(2,38)=8.24, p=0.001). Moreover, similar to our previous work using the same functional experiment (Grotheer et al., 2018), the predicted VWFA-2 showed a task by stimulus interaction (repeated measures ANOVA: F(2,38)=5.93, p=0.006). The response characteristics of the predicted VWFA-2 fROI hence closely match those observed for measured VWFA-2 fROIs.

Overall, our data show that the endpoints of the ILF, AF, and VOF in combination with cortical T_1_ predict reading-related responses across lateral VTC as well as the specific location of the VWFA-2.

## 4. Discussion

In the current study we used a multimodal approach to test the hypothesis that white matter fascicles and gray matter microstructure covary with reading-related responses in the lateral ventral temporal cortex (lateral VTC). We find that i) the endpoint densities of the arcuate fasciculus (AF), inferior longitudinal fasciculus (ILF), and vertical occipital fasciculus (VOF) predict reading-related responses, ii) gray matter microstructure, as assessed by proton relaxation time, T_1_, further improves the prediction of reading related-responses, compared to using only information from endpoints of white matter fascicles, and iii) a model that combines the endpoints of the AF, ILF, and VOF with cortical T_1_ accurately predicts the map of reading-related responses in lateral VTC as well as the location of the visual word form area, a category-selective region that plays a critical role in reading.

Previous research has provided compelling evidence for the role of white matter connections in driving the consistent organization of functional regions in VTC (Bi et al., 2015; Osher et al., 2016; Saygin et al., 2016, 2012), including the development of reading-related responses during childhood (Saygin et al., 2016). Most previous studies relied on estimating the “white matter fingerprint”, an approach that utilizes brain parcellations (e.g., anatomical parcels from FreeSurfer) and measures the amount of pairwise white matter tracts between each functional voxel in one parcel to all other cortical parcels. A weighted sum of these pairwise connections is used as a predictor for that voxel’s functional responses (Osher et al., 2016; Saygin et al., 2016, 2012). While the prior approach has been important for showing relationships between white matter tracts and functional responses, it did not reveal whether there is a correspondence between specific, known white matter fascicles and the spatial layout of reading-related responses in lateral VTC. By developing a new approach that uses well-established components of the human brain’s white matter anatomy, specifically, the major white matter fascicles of the brain, as predictors of functional responses, the current study elucidates the systematic coupling between functional activations and white matter fascicles for the first time.

We found that three fascicles of the brain, the AF, ILF, and VOF, predict the spatial layout of reading-related responses in lateral VTC, suggesting that these fascicles form the backbone of the reading-network. Diffusion properties of these fascicles have previously been correlated with people’s performance in reading tasks, supporting the notion that they play a critical role in connecting different parts of the reading network. For instance, fractional anisotropy (FA) in the left AF is related to reading ability during childhood development (Wang et al., 2017) and correlates with phonological awareness in both typical (Yeatman et al., 2011) and impaired (Su et al., 2018; Vandermosten et al., 2012) readers. Similarly, atypical development of FA in the ILF is associated with poor reading proficiency (Su et al., 2018; Yeatman et al., 2012a), while lesions to the ILF have been associated with pure alexia (Epelbaum et al., 2008). Finally, the VOF has been re-discovered recently (Takemura et al., 2016; Weiner et al., 2016; Yeatman et al., 2014b), is proposed to carry top-down feedback signals during reading (Kay and Yeatman, 2017), and diffusion measures in this tract have also been linked to early literacy skills in children (Broce et al., 2019).

Interestingly, our data shows that even though the AF, ILF, and VOF are all predictive of the spatial layout of reading-related responses, high endpoint densities of the AF and ILF predicts vertices showing high responses during reading, whereas high endpoint density of the VOF instead predicts low responses during reading. While previous studies agree that the AF and ILF connect to the VWFA (Bouhali et al., 2014; Grotheer et al., 2019; Yeatman et al., 2013), a region in VTC that shows a preference for reading, there have been conflicting results on whether or not the VOF connects to this region (Bouhali et al., 2014; Grotheer et al., 2019; Yeatman et al., 2013). These conflicting results can likely be explained by the fact that the VWFA is divided into two distinct regions, VWFA-1 and VWFA-2, also referred to as pOTS-words and mOTS-words, respectively, which show different connectivity to the VOF (Lerma-Usabiaga et al., 2018). Here we focused only on the VWFA-2, as VWFA-1 could not be identified reliably when comparing responses in a reading task with responses during other tasks performed on identical visual stimuli. This, in turn, matches findings that the VWFA-1 is a more “perceptual” visually-driven region whereas the VWFA-2 is a higher-level region involved in lexical / semantic processing (Lerma-Usabiaga et al., 2018) as well as the more general proposal of an posterior-to-anterior gradient in processing level related to reading along the lateral VTC (Taylor et al., 2019; Vinckier et al., 2007). While the VWFA-1 is suggested to connect to the VOF, the more anterior VWFA-2, which was investigated in the current study, is proposed not to connect to the VOF (Lerma-Usabiaga et al., 2018), which matches our findings of low reading-related responses in vertices that have a high endpoint density of the VOF.

The current work has implications for our understanding of the neural underpinnings of reading. Given that the ILF, AF, and VOF are predictive of the spatial layout of reading-related responses, and white matter structure constraints reading-related responses during development (Saygin et al., 2016), these fascicles likely play a causal role in scaffolding the neural architecture required for reading. Future studies in atypical populations, for instance those evaluating children with dyslexia (e.g. Kraft et al., 2016; Niogi and McCandliss, 2006; Vanderauwera et al., 2017; Yeatman et al., 2012a; Zhao et al., 2016), could test if structural abnormalities in these fascicles precede and predict difficulties in reading acquisition and whether these abnormalities may result in subtle differences in the spatial layout of reading-related responses in lateral VTC (e.g. Kubota et al., 2019). Another interesting direction for future research would be to probe if short-range white matter connections (e.g. Gomez et al., 2015), which are not considered here, also contribute to the spatial layout of reading-related responses in VTC. Rather than evaluating the entire white matter connectome, as in the fingerprinting approach, we would suggest a more targeted approach that evaluates the contribution of pairwise connections between functional regions of interest (Grotheer et al., 2019), as such data would be more easily interpretable.

In the current study, in addition to investigating the role of the white matter, we also began to explore the additional hypothesis that the local gray matter tissue properties contribute to the consistent spatial layout of reading-related response in lateral VTC. The notion that differences in microstructure across cortical regions may co-localize with different functional regions dates back to the very birth of neuroscience (Brodmann, 1909). Recent methodological improvements, particularly the development of quantitative MRI (Lutti et al., 2014; Mezer et al., 2013), now enable us to test this prediction *in vivo*, for the first time. We found that proton relaxation time (T_1_) at each vertex, measured with qMRI, improves the prediction of reading-related responses compared to a model that relies only on white matter fascicles. This aligns well with previous work showing that i) VTC contains several distinct cytoarchitectonic regions (Caspers et al., 2013; Lorenz et al., 2017), iii) functional regions in VTC co-localize with cytoarchitectonic regions (Weiner et al., 2017), and iii) T_1_ varies between different functional regions in VTC (Gomez et al., 2017; Natu et al., 2019). However, it has also been shown that T_1_ changes during development and correlates with task performance (Gomez et al., 2017). Thus, an open question is whether T_1_ development precedes reading acquisition and the emergence of reading-related functional activations, as is the case for white matter connectivity (Saygin et al., 2016), or whether reading acquisition instead changes the local tissue microstructure, for instance due to tissue proliferation. In the latter case, the predictiveness of T_1_ would be a consequence rather than a cause of the consistent organization of lateral VTC. The current work in adults should be considered a proof-of-concept that gray matter tissue structure can predict functional responses in VTC and future developmental work can address if gray matter structure constrains the organization of VTC during development. Such studies could go beyond T_1_ measurements and probe the predictive value of various different aspects of the gray matter structure that can be assessed *in vivo*, including both gray matter microstructure, such as myelination (Lutti et al., 2014; Natu et al., 2019; for review see Edwards et al., 2018; Weiskopf et al., 2015), as well as, macrostructure, such as cortical folding (Weiner et al., 2018, 2014) and cortical thickness (Natu et al., 2019; Sowell et al., 2004, 2003).

In conclusion, the current study showed that cortical terminations of three key fascicles, the AF, ILF, and VOF, in combination with gray matter microstructure predicts the map of reading-related responses in lateral VTC as well as the location of the visual word form area. The current study deepens our understanding of the neural substrates of reading and opens a new avenue of research that carefully assesses the role of various features of the gray and white matter anatomy in driving the organization of the VTC and the brain.

## Acknowledgements

This research was supported by the National Institute of Health (NIH; grant 1R01EY023915 awarded to KGS as well as grants 1RF1MH121868 and R01HD09586101 awarded to JDY), by the Deutsche Forschungsgemeinschaft (DFG; grant GR 4850/1-1 awarded to MG) and by an Innovation Grant from the Stanford Center for Cognitive and Neurobiological Imaging (CNI; awarded to MG).

## Author Contribution

MG collected the data and developed code used for data analyses. MG, KGS and JDY analyzed the data. MG, KGS and JDY wrote the manuscript.

## Competing Interests

The authors declare no competing interests.

## Notes

### Competing Interest Statement

The authors have declared no competing interest.

